# GraphMana: graph-native data management for population genomics projects

**DOI:** 10.64898/2026.04.11.717925

**Authors:** Ehsan Estaji, Shi-Wei Zhao, Zhao-Yang Chen, Shuai Nie, Jian-Feng Mao

**Affiliations:** Umeå Plant Science Centre, Department of Plant Physiology, Umeå University, SE-90187, Umeå, Sweden; Rice Research Institute, Key Laboratory of Genetics and Breeding of High Quality Rice in Southern China (Co-construction by Ministry and Province), Ministry of Agriculture and Rural Affairs, Guangdong Key Laboratory of New Technology in Rice Breeding, Guangdong Academy of Agricultural Sciences, 510640, Guangzhou, China

**Author notes:** Contributing authors. These authors contributed equally to this work (Ehsan Estaji, Shi-Wei Zhao).

**Keywords:** population genomics, data management, graph database, variant call format, genotype storage, reproducibility

## Abstract

Population genomics projects rely on fragmented file-based workflows that lose provenance and require full reprocessing when samples are added. Graph-Mana stores variant data in a graph database as packed genotype arrays with pre-computed population statistics, enabling incremental sample addition, provenance tracking, cohort management, and export to 17 formats. Two access paths serve different needs: a FAST PATH reading population-level arrays in ***O*(*K*)** time and a FULL PATH unpacking per-sample genotypes in ***O*(*N*)** time. On human 1000 Genomes data (3,202 samples, 70.7M variants), Graph-Mana completed a 46-operation lifecycle in 98 minutes from a single persistent database.

---

Population genomics projects at the scale of hundreds to tens of thousands of samples face a data management problem that existing tools do not natively integrate. Adding new sequencing batches forces regeneration of every downstream file—every VCF [9], PLINK binary [5], TreeMix input [12], and site frequency spectrum [10]— because flat-file formats encode the complete sample set and cannot be extended in place. Workflow managers such as Snakemake [17] and Nextflow can automate regeneration, but they coordinate files externally rather than storing data, provenance, and annotations in a single queryable structure. Database-backed systems exist: Gemini [16] stores variants in SQLite for annotation-driven queries but does not manage multi-format export or incremental sample addition; TileDB-VCF provides columnar variant storage with sample-level append but lacks built-in population statistics and export format diversity; Hail [4] provides scalable matrix computation but treats annotations and provenance as external metadata and targets biobank-scale rather than mid-scale project management. This coordination burden falls in a gap between single-investigator work (where manual tracking suffices) and biobank-scale programs.

Even below biobank scale, the problem is growing. Projects such as the 3,000 Rice Genomes, 1000 Bull Genomes, and regional human sequencing initiatives routinely operate in the hundreds-to-tens-of-thousands range where sample counts increase with each sequencing batch, analytical scope broadens, and multi-format data sharing intensifies—yet the underlying file-based infrastructure remains unchanged.

These coordination failures compound through the weekly rhythm of a working project (Supplementary Figs. 1 and 2). A sequencing core delivers 200 new samples, triggering regeneration of every VCF, PLINK binary, and allele frequency table. A collaborator requests an EIGENSTRAT [13] file for a specific population subset, filtered by MAF; producing it requires a one-off script whose parameters go unrecorded. A ClinVar update arrives, but because annotations live inside the VCF header, incorporating new functional data means rewriting the entire genotype file. When questions arise about which samples or filters produced a particular result shared months earlier, the only recourse is to reconstruct the answer from directory timestamps and notebook entries. None of these tasks is computationally hard—bcftools [6] merges files in minutes, PLINK recodes in seconds—but the accumulation of untracked conversions, orphaned scripts, and stale file copies creates a coordination overhead that grows with each new collaborator, annotation source, and export format.

The root cause is structural: population genomics data is inherently relational— variants belong to chromosomes, samples to populations, genes carry functional annotations with versioned provenance—but flat files flatten these relationships into disconnected tables. Several architectures could address this (Table 1). Relational databases require JOIN operations and lack native support for extensible byte-array properties. Columnar stores and Hail excel at matrix computation but treat annotations and provenance as external metadata. Graph databases store data as *nodes* connected by typed *edges*, and queries traverse these connections directly, making lifecycle operations—incremental addition, annotation versioning, cohort definition, provenance binding—persistent and local rather than requiring full-dataset regeneration. Other database models could also address parts of this problem; the graph-native model provides a particularly natural fit because the relationships in population genomics data (variants to chromosomes, samples to populations, genes to pathways) are numerous, typed, and frequently traversed.

**Table 1.** Data management architectures for population genomics. Operations natively supported (•), supported only with external scripting (∘), or not supported (—). The five columns compare flat files (VCF/BCF, PLINK, EIGENSTRAT, etc.), relational databases, columnar stores (TileDB-VCF), Hail-style matrix frameworks, and GraphMana. GraphMana is the only architecture that natively supports all six lifecycle operations: incremental sample addition, annotation versioning, provenance binding, cohort-as-query, multi-format export, and in-place annotation update.

| Operation | Flat files | Relational DB | TileDB-VCF | Hail | GraphMana |
| --- | --- | --- | --- | --- | --- |
| Incremental sample addition | — | ○ | ● | ○ | ● |
| Annotation versioning | — | ○ | — | — | ● |
| Provenance binding | — | ○ | — | ○ | ● |
| Cohort as query | — | ● | ○ | ● | ● |
| Multi-format export (17) | ○ | — | — | ○ | ● |
| In-place annotation update | — | ● | — | — | ● |

GraphMana implements this graph-native approach as what we term a *persistent analytical record* : a single queryable database in which genotype data, population statistics, annotations, cohort definitions, and provenance coexist and evolve together (Fig. 1a; Supplementary Fig. 3). Each biallelic variant is a node carrying a packed genotype array (2 bits per sample, 4 per byte) alongside pre-computed population-level allele counts, sample counts, and frequency arrays (Fig. 1b). This encoding reduces storage 125-fold compared with per-sample graph edges—the naive model a graph database designer would first consider—and the encoding density is comparable to matrix-based stores such as Hail and TileDB-VCF, while retaining the typed-edge semantics that make lifecycle operations local (Supplementary Table 1). The packed representation creates a two-tier access model. Most population genetics queries—TreeMix [12], site frequency spectra [10, 11], allele frequency tables—need only population-level aggregates, which are *K*-element arrays constant in size regardless of sample count; GraphMana serves these via a FAST PATH in *O*(*K*) time (Fig. 1c; Supplementary Fig. 4). Per-sample formats (VCF, PLINK [5], EIGENSTRAT [13], Beagle [14], STRUCTURE [15], and 11 others) use a FULL PATH that unpacks genotypes in *O*(*N*) time (Supplementary Fig. 5). The practical consequence is that as projects grow from 1,000 to 50,000 samples, the most common analytical exports remain instantaneous (Supplementary Table 2). In total, 17 formats are supported (Supplementary Note 1), of which 6 have been validated against downstream tools and 6 against format specifications (Supplementary Tables 3–4). Genotype roundtrip fidelity exceeds 99.999%, with the residual mismatches confined to multi-allelic position ambiguity rather than data loss (Supplementary Tables 5–6).

**Figure 1.**
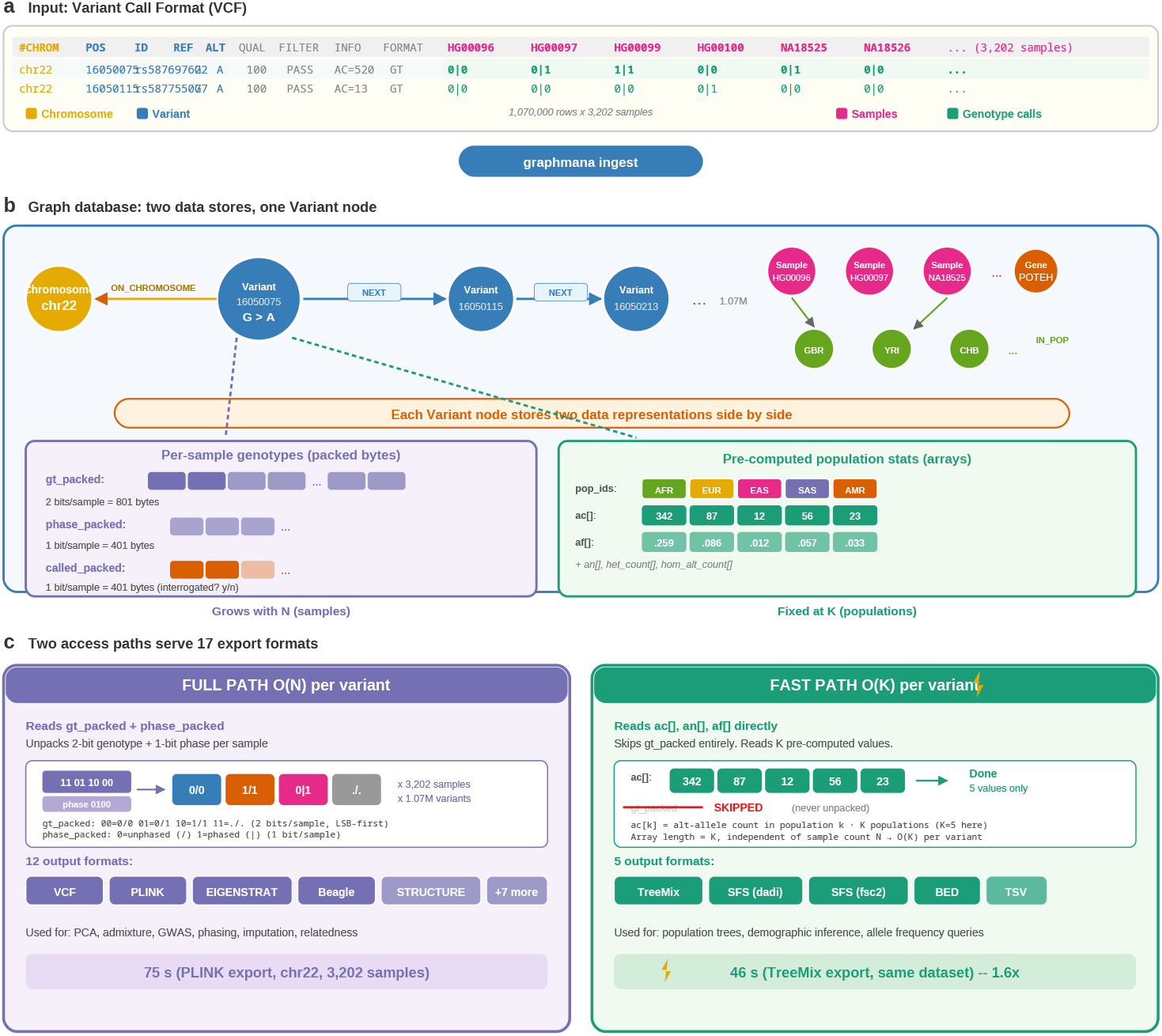
From VCF to graph database to 17 export formats. (**a**) Genetic variants in a flat file (Variant Call Format, VCF), with columns color-coded by their destination in the graph—chromosome (gold), variant identity (blue), sample identifiers (pink), and per-sample genotype calls (green). (**b**) After `graphmana ingest`, the same genetic variants are stored in a graph database: Variant nodes are linked in genomic order by NEXT relationships, attached to Chromosome nodes via ON CHROMOSOME, and associated with Sample, Population, and Gene nodes via typed edges. Each Variant node stores two data representations side by side, one feeding each of the two access paths shown in panel c. Left (purple, FULL PATH source): per-sample genotypes encoded as three parallel packed byte arrays— `gt_packed` (2 bits per sample; 801 bytes for 3,202 samples), `phase_packed` (1 bit per sample), and `called_packed` (1 bit per sample, flagging whether each sample was actually interrogated at this site). All three arrays grow linearly with *N*. Right (green, FAST PATH source): pre-computed population arrays (`pop_ids[], ac[], af[], an[], het_count[], hom_alt_count[]`) containing *K* elements (*K* = number of populations), constant in size regardless of *N*. Per-population denominators are computed against `called_packed`, so allele frequencies remain statistically honest even when incremental batches carry different site lists. This dual representation is the architectural foundation for the two export paths. (**c**) The `graphmana export` command selects one of two access paths based on the requested format. FULL PATH (purple, *O*(*N*) per variant) unpacks `gt_packed` and `phase_packed` and translates to 12 format-specific encodings (VCF, PLINK, EIGENSTRAT, Beagle, STRUCTURE, and seven others); used for PCA, admixture, GWAS, phasing, imputation, and relatedness (75 s for PLINK export of chr22, 3,202 samples). The unpacking strip illustrates a four-sample example: `gt_packed` byte `11 01 10 00` (read LSB-first, so the rightmost two bits are sample 1) decodes to `0/0, 1/1, 0/1, ./.` for samples 1–4; the paired `phase_packed` bits `0100` mark sample 3 as phased, rendering its heterozygous call as `0|1`. The 2-bit genotype code is `00`=HomRef, `01`=Het, `10`=HomAlt, `11`=Missing; the 1-bit phase code is `0`=unphased (/) and 1=phased (|). Multi-allelic sites are split into biallelic Variant nodes during `graphmana_ingest` (bcftools-style normalization), so each node always carries a biallelic 2-bit code; a `1/2` input genotype is stored as Het at both resulting Variant nodes. FAST PATH (green, *O*(*K*) per variant) reads `ac[], an[], af[]` directly, skipping `gt_packed`; the per-population allele-count array has length *K* (number of populations, *K*=5 in the illustration) and is constant in size regardless of *N*. Used for population tree inference, demographic modeling, and frequency queries via 5 formats (TreeMix, SFS-dadi, SFS-fsc2, BED, TSV; 46 s for TreeMix on the same dataset; the modest 1.6× speedup at chromosome scale reflects Neo4j query overhead, which diminishes at whole-genome scale where the per-variant *O*(*K*) vs *O*(*N*) cost dominates). FAST PATH runtime is independent of sample count.

Because relationships are explicit graph edges rather than columns embedded in flat files, annotations can be updated by modifying edge properties without touching genotype data—achieving a 27-fold speedup over full VCF rewrites (3.5 s versus 96 s for 53,000 regulatory regions; Fig. 2b). Cohorts are defined as graph queries rather than file extractions. Every export generates a machine-readable manifest recording the software version, filters, and sample set, so that provenance questions are answered by querying metadata rather than reconstructing it from file timestamps (Supplementary Fig. 6).

**Figure 2.**
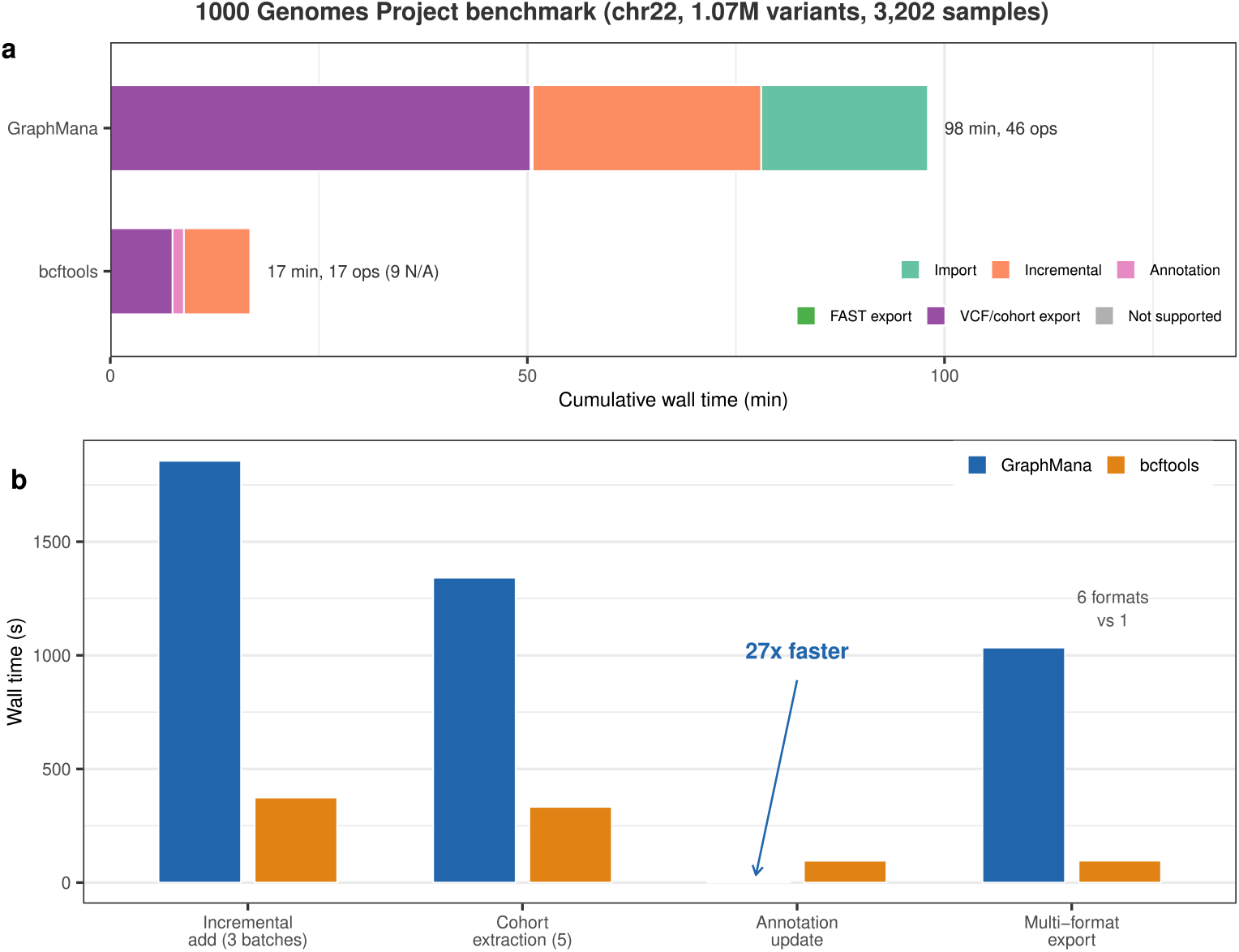
**Human 1000 Genomes Project benchmark** (chromosome 22, 1.07 million variants, 3,202 samples). (**a**) Lifecycle simulation across 46 operations spanning import, filtering, cohort extraction, annotation update, and multi-format export. GraphMana completes all 46 operations from a single persistent database; bcftools completes 17 of 26 attempted operations, with 9 operations (multi-format export, in-place annotation update, cohort management) having no equivalent in a flat-file workflow and shown in grey. (**b**) Per-task timing comparison for the operations supported by both tools. Annotation updates are 27× faster in GraphMana (3.5 s versus 96 s for 53,000 regulatory regions) via in-place edge modification. Multi-format export covers 6 formats (VCF, PLINK, EIGENSTRAT, TreeMix, SFS, BED) versus VCF only for bcftools. Per-task timings favor bcftools for shared file-streaming operations (3–5× faster); GraphMana’s advantage emerges over the project lifecycle by eliminating repeated file regeneration and manual coordination.

Incremental sample addition extends packed arrays without modifying existing data. On the human 1000 Genomes Project [3] (70.7 million variants, 3,202 whole-genome samples), adding 234 samples took 182 minutes via a CSV-to-CSV rebuild; approximately 95% of variants required only zero-byte extension (Supplementary Table 7). No downstream result was invalidated because all queries operate on the live database.

Incremental addition from independently called VCFs introduces a correctness hazard: positions absent from a batch are ambiguous between “called HomRef” and “not interrogated.” GraphMana resolves this by storing a per-sample `called_packed` bit on every Variant node; allele-frequency denominators exclude not-interrogated samples, keeping estimates honest under batch-by-batch growth. The recommended upstream workflow is joint calling from gVCFs via GenomicsDB or GLnexus (Supplementary Note 5; Supplementary Fig. 8).

We benchmarked GraphMana against bcftools [6] on chromosome 22 (1.07 million variants, 3,202 samples) across five tasks (Supplementary Tables 8–10), with whole-genome export timings reported separately (Supplementary Table 11). Graph-Mana completed all 46 lifecycle operations in 98 minutes (97.6 and 98.4 across two runs); bcftools completed 17 of 26 in 17 minutes, with 9 operations—multi-format export, in-place annotation, cohort management—having no equivalent (Fig. 2a). For shared operations, bcftools was 3–5× faster per task (Fig. 2b), reflecting sequential file-streaming efficiency versus graph query overhead. The difference is architectural: bcftools optimizes individual file operations, whereas GraphMana eliminates the repeated regeneration and manual coordination between them. We note that bcftools is a file-manipulation tool, not a database; Table 1 provides a conceptual feature comparison against database-backed alternatives. The primary advantage of the FAST/FULL PATH distinction is not raw per-operation speed (1.6× at chromosome scale, where Neo4j query overhead dominates) but the ability to serve 17 export formats from a single persistent store without re-exporting from flat files. At whole-genome scale (85M variants), the per-variant *O*(*K*) versus *O*(*N*) difference becomes the dominant term.

GraphMana currently represents variants as biallelic nodes; multi-allelic records are decomposed at import and reconstructed during VCF export (Supplementary Note 2). Structural variants are stored with typed properties. gVCF input is supported by joint-calling upstream (GenomicsDB or GLnexus) and ingesting the resulting multi-sample VCF; raw per-sample gVCF reference blocks are not parsed directly by GraphMana, but the HomRef-vs-Missing distinction they encode is preserved by `called_packed` (Supplementary Note 5 and Supplementary Fig. 8). The platform operates as a single-writer system with multi-reader support via replicas.

Packed encoding scales linearly: ⌈*N*/4⌉ bytes per variant for genotypes, ⌈*N*/8⌉ for phase. All scaling estimates assume human whole-genome data (~ 85M biallelic variants); for other species or designs, requirements scale proportionally by variant count (Supplementary Tables 1–2). At 100–10,000 samples, all operations are interactive; beyond 50,000 samples, the single-node architecture becomes a bottleneck, and distributed frameworks [4] are more appropriate. The current implementation uses Neo4j Community Edition [2]; the design is not vendor-specific. The companion GraphPop engine [1] provides graph-native population genetics computation on the same database.

GraphMana has several limitations. The single-node Neo4j architecture imposes a practical ceiling of approximately 50,000 whole-genome samples; beyond this scale, distributed frameworks such as Hail are more appropriate. The Neo4j data directory must reside on local SSD—network filesystems cause severe performance degradation, limiting cloud and shared-storage deployments. The system operates as a single-writer with multi-reader support; concurrent write transactions are not supported. Multi-allelic variants are decomposed into biallelic nodes during import and reconstructed at VCF export; users of tools requiring intact multi-allelic records should verify roundtrip fidelity. Raw per-sample gVCF reference blocks are not parsed directly; joint calling upstream is required for correct HomRef-vs-Missing semantics.

Despite these constraints, for projects in the 100–50,000-sample range that constitute the majority of population genomics studies, GraphMana replaces ephemeral file-based workflows with a persistent analytical record that addresses a reproducibility gap individual tool improvements cannot close. Software, documentation (12 vignettes, 67 command reference pages, of which 58 document primary commands and 9 document subcommands), and pre-built databases are available under the MIT license. All data and code follow FAIR principles: the software is indexed on PyPI, source is on GitHub, and benchmark data and pre-built databases are deposited on Zenodo with persistent DOIs.

## Online Methods

### Graph database technology

A graph database stores data as nodes (entities) and edges (typed, directed relationships), with key-value properties on both. Unlike relational databases, which reconstruct relationships through table joins at query time, graph databases traverse pre-materialized connections in constant time per hop—a property known as index-free adjacency. This distinction matters for genomic data because the relationships between variants, chromosomes, genes, populations, and annotation sources are numerous, typed, and frequently traversed.

In GraphMana’s property-graph model, a Variant node connects to its Chromosome via an ON_CHROMOSOME edge, to Gene nodes via HAS_CONSEQUENCE edges carrying annotation properties (consequence type, impact level, annotation version), and to the ordered variant chain via NEXT edges. Sample nodes connect to Population nodes via IN_POPULATION edges. Updating an annotation version modifies HAS_CONSEQUENCE edge properties without touching genotype arrays; defining a cohort is a graph traversal that produces a sample list without extracting files; and provenance is recorded as IngestionLog nodes linked to the operations that produced them. These operations are local to the edges or nodes involved, not to the entire dataset.

We chose Neo4j Community Edition [2] for its mature property-graph implementation, the Cypher query language, ACID transaction support, and free open-source licensing. The graph-native design principles—typed nodes and edges, byte-array properties on nodes, edge-independent updates—are not specific to Neo4j and could be implemented on other property-graph databases. A concrete end-to-end project workflow is presented in Supplementary Note 3, and Supplementary Note 4 provides a detailed introduction for readers unfamiliar with graph database concepts.

### Software architecture

GraphMana comprises a Python command-line application (~ 21,000 lines of source code at the time of submission) with a bundled Java plugin (~ 2,900 lines) for server-side Neo4j procedures. The Python layer handles VCF parsing via cyvcf2 [8], genotype packing and unpacking via NumPy (complementing scikit-allel [7] for lower-level variant access), CSV generation for bulk database loading, and all 17 export format writers. The Java plugin provides optional server-side packed array extension for incremental imports, avoiding Bolt protocol round-trips for large batch operations.

The 58 CLI commands are organized into nine functional domains: data import and integration (5 commands), annotation management (9), data export (2), sample and cohort management (11), quality control and verification (3), provenance and state tracking (7), database administration (8), status and reporting (5), and infrastructure and deployment (6). The full command hierarchy is shown in Supplementary Fig. 7. A companion Python package (graphmana-py) provides a Jupyter-compatible API returning pandas DataFrames for interactive analysis. The test suite comprises 1,451 unit and integration tests covering genotype packing, phase conventions, byte boundary conditions, export format contracts, filter chains, and database operations.

### Data encoding

The graph database contains five primary node types—Variant, Sample, Population, Chromosome, and Gene—connected by four relationship types: ON_CHROMOSOME, IN_POPULATION, HAS_CONSEQUENCE, and NEXT (chromosomal ordering). Variant nodes store genotype data as packed byte arrays using a 2-bit encoding: gt_packed stores diploid genotypes (00 = homozygous reference, 01 = heterozygous, 10 = homozygous alternate, 11 = missing), yielding 4 genotypes per byte. Phase information is stored separately in phase packed (1 bit per sample, 8 per byte), and ploidy status in ploidy packed (1 bit per sample; null when all samples are diploid). Genotype values are remapped from cyvcf2’s internal coding (where homozygous alternate = 3 and missing = 2) to the sequential 2-bit encoding at import time; the remap is applied exactly once per code path and verified by 33 dedicated unit tests. This packed representation achieves a 125-fold storage reduction compared with an alternative design using individual Sample-to-Variant edges (Supplementary Table 1).

Each Variant node also carries *K*-element population-level pre-computed arrays: pop_ids (population identifiers), ac (allele counts per population), an (allele numbers), af (allele frequencies), het count, hom_alt_count, and het_exp (expected heterozygosity under Hardy–Weinberg equilibrium). These arrays have *K* elements where *K* is the number of populations (typically 5–30), making them constant in size regardless of sample count *N*. This property enables the FAST PATH access pattern: queries that need only population-level statistics (TreeMix, SFS, allele frequency tables) read these pre-computed arrays directly without unpacking any per-sample genotype data. Full encoding details and a worked multi-allelic decomposition example are provided in Supplementary Note 2.

### Import pipeline

GraphMana uses a two-step split pipeline designed for both workstations and HPC clusters. The first step (prepare-csv) parses VCF files using cyvcf2 for streaming access, packs genotypes into byte arrays, computes population statistics from the genotype calls, and writes Neo4j-compatible CSV files. This step is embarrassingly parallel across chromosomes and requires no running database instance, making it suitable for batch job submission on compute nodes without Neo4j. The second step (load-csv) uses neo4j-admin import for bulk loading, which writes directly to the Neo4j store files and bypasses the transaction engine, achieving import speeds limited only by disk I/O.

On our benchmark hardware (32 CPU cores, 64 GB RAM, NVMe SSD), CSV generation for 2,500 human whole-genome samples across 22 chromosomes (70.7 million variants) completed in 95 minutes with 16 parallel threads (single-thread estimate: ~25 hours, extrapolated from per-chromosome timings); bulk loading of the resulting 214 GB of CSV files completed in 3 minutes.

### Incremental sample addition

Adding new samples to an existing database extends the packed genotype arrays of all existing Variant nodes. GraphMana provides three strategies of increasing throughput: (1) Cypher transactions on a running database instance, suitable for small additions (fewer than 10,000 variants); (2) an export-extend-reimport path that reads variant data from Neo4j, extends arrays in Python, and reimports via neo4j-admin; and (3) a CSV-to-CSV rebuild that reads the existing CSV checkpoint directly at NVMe speed, extends arrays using NumPy vectorized operations, and produces a new CSV set for reimport. The CSV-to-CSV path is fastest for whole-genome databases: on our benchmark dataset, extending 70.7 million variants with 234 new samples completed in 182 minutes, of which 160 minutes was sequential CSV I/O and array extension, 15 minutes was neo4j-admin import, and 7 minutes was VCF parsing and database restart. Approximately 95% of variants (69.6 million of 70.7 million) were on chromosomes absent from the new VCF and required only HomRef extension—appending zero bytes to packed arrays without unpacking and repacking.

### Correctness validation

We validated genotype fidelity by importing human 1000 Genomes Project chromosome 22 data (897,645 biallelic SNPs, 5 samples), exporting to phased VCF, and comparing per-sample genotypes with the original using bcftools. Concordance exceeded 99.999%, with 2–8 mismatches per sample confined to multi-allelic positions where position-based joining during comparison could not distinguish co-located biallelic records. A systematic correctness matrix covering biallelic SNPs, indels, multiallelic sites, structural variants, phased and unphased genotypes, missing data, haploid and diploid ploidy, and annotation-layer independence is provided in Supplementary Tables 5–6.

### Benchmark methodology

All benchmarks used the human 1000 Genomes Project high-coverage dataset [3] (GRCh38). Chromosome 22 (1,066,557 variants, 3,202 samples, 26 populations) served as the primary target for the full comparison matrix. The 3,202 samples were split deterministically (seed = 42) into a base cohort of 2,500 and three incremental batches of 234 each.

We compared GraphMana against bcftools [6] (version 1.17) as the single comparison target, chosen as the stronger baseline for VCF-centric operations. Five benchmarks were conducted: (1) incremental sample addition (GraphMana CSV-to-CSV rebuild versus bcftools merge, 3 rounds); (2) cohort-specific VCF extraction (5 superpopulation cohorts); (3) multi-format export (6 formats from a single source; bcftools supports VCF only); (4) annotation update (53,000 BED regulatory regions; in-place edge modification versus VCF rewrite); and (5) a lifecycle simulation comprising 7 phases of interleaved imports, exports, and annotation updates. All timings report wall-clock time on a workstation with 32 CPU cores, 64 GB RAM, and NVMe SSD. The graph database was configured with 16 GB heap and 16 GB page cache. Each benchmark was run twice; reported values are from the second run (warm page cache), with both runs reported for the lifecycle total. Export timings for PLINK and TreeMix were additionally measured in triplicate; we report the median (PLINK: 75 s, range 52–75, lower bound reflects warm page cache; TreeMix: 46 s, range 46–47). We emphasize that per-operation timings favor bcftools, which is optimized for sequential file streaming; GraphMana’s advantage lies in avoiding repeated file regeneration across the project lifecycle. Benchmark scripts and raw timing data are deposited at Zenodo. The benchmark hardware (32 cores, 64 GB RAM, NVMe SSD) is high-end; on lower-specification machines, absolute timings will be proportionally slower, but the lifecycle advantage—fewer operations rather than faster operations—is hardware-independent.

### Scalability

Packed encoding scales linearly: storage per variant is ⌈*N*/4⌉ bytes for genotypes and ⌈*N*/8⌉ bytes for phase, where *N* is the number of samples. All scaling estimates in this work assume human whole-genome data (~85 million biallelic variants per genome); for species or designs with fewer variants (exome sequencing at ~5M variants, rice at ~29M), storage and time requirements scale proportionally by variant count. At 100– 10,000 human whole-genome samples, all operations are interactive. At 10,000–50,000 samples, FAST PATH operations remain instantaneous because population arrays are constant size regardless of *N*, but FULL PATH exports scale linearly and become the bottleneck. Beyond 50,000 samples, the single-node graph database architecture imposes practical limits due to per-variant query latency and transaction log pressure. At biobank scale (500,000+ samples), distributed computing frameworks such as Hail [4] are more appropriate. Supplementary Tables 1–2 provide detailed storage and performance estimates from 3,202 to 500,000 samples.

### Deployment

GraphMana installs entirely in user space without administrator privileges. The primary installation method is `pip install graphmana`, which bundles the pre-built Java procedures JAR as package data. The `graphmana setup-neo4j` command downloads and configures Neo4j Community Edition, automatically deploys the JAR to the plugins directory, and optionally downloads Eclipse Temurin JDK 21 via the `--install-java` flag for systems without Java. For HPC clusters, a two-step split pipeline separates CSV generation (which runs on any compute node without Neo4j) from bulk loading (which requires the database host). SLURM and PBS job scripts are provided. The Neo4j data directory must reside on local SSD or scratch storage; network filesystems (NFS, Lustre, GPFS) cause severe performance degradation, and GraphMana includes a filesystem check command that warns users before import.

## Supporting information

all supplementary materials

## Data availability

GraphMana is available at https://github.com/jfmao/GraphMana under the MIT license. Neo4j Community Edition, used as the underlying database engine, is distributed under the GPL v3 license. Benchmark data and pre-built databases are deposited at Zenodo (https://doi.org/10.5281/zenodo.19472835).

## Code availability

All source code is at https://github.com/jfmao/GraphMana under the MIT license.

## Acknowledgements

This work was partially supported by the Wallenberg Initiatives in Forest Research (WIFORCE) funded by the Knut and Alice Wallenberg Foundation. Genomic data processing and analyses were performed using resources provided by the Swedish National Infrastructure for Computing (SNIC), through the High Performance Computing Centre North (HPC2N) at Umeå University. The human 1000 Genomes Project dataset [3] was used under its open data access terms.

## Author contributions

Conceptualization and design: J.-F.M., E.E., and S.N. Implementation and testing: E.E., S.-W.Z., Z.-Y.C., and S.N. Writing and editing: J.-F.M., E.E., and S.-W.Z.

