## Supplementary material for "GraphMana: graph-native data management for population genomics projects": all supplementary materials

#### Contents

**1 Supplementary Figure Legends** **2**

**2 Supplementary Table Legends** **5**

**3 Supplementary Note 1: Export Format Reference** **7**

**4 Supplementary Note 2: Variant Representation and Data Encoding** **9**

**5 Supplementary Note 3: Concrete Workflow Example** **11**

**6 Supplementary Note 4: Graph Database Concepts for Non-Specialists** **13**

**7 Supplementary Note 5: Preserving the HomRef-vs-Missing Distinction Across Incremental Batches** **15**

**8 Supplementary References** **18**

**9 Supplementary Figures** **19**

**10 Supplementary Tables** **28**

### 1 Supplementary Figure Legends

**Supplementary Figure 1. A population genomics data manager's year — conventional file-based workflow.** Timeline of recurring data management activities in a typical population genomics project using conventional file-based workflows. Seven phases: (1) reception of the first VCF batch with QC, filtering, and population assignment; (2) functional and clinical annotation; (3) format conversion using separate scripts per downstream tool; (4) population genetics analyses and summary statistics; (5) arrival of a second batch triggering a complete restart—re-merge, re-QC, re-filter, re-annotate, re-convert all formats, and re-run all analyses (2–4 weeks of lost work, repeated with every subsequent batch); (6) ongoing requests including collaborator subsets, reviewer re-analyses, annotation updates, liftover, and additional batches 3–5; (7) provenance failures—inability to answer which VCF was used for which analysis, what filters were applied, or which annotation version was current. Over a typical project lifetime this workflow involves 5–15 merge cycles, 50+ format conversions, 200+ intermediate files, and no systematic provenance tracking.

**Supplementary Figure 2. A population genomics data manager's year — the same project managed through GraphMana.** The same year as Supplementary Figure 1, now managed through GraphMana. Phase 1 (minutes, not weeks): a single `graphmana ingest` command applies filters, packs genotypes, pre-computes population statistics, and builds the graph. Phase 2 (seconds): `graphmana annotate` adds functional and clinical annotations as versioned graph edges in-place. Phase 3 (seconds per export): any of 17 output formats via a single `graphmana export` command with filters as flags—no conversion scripts. Every export auto-generates a `.manifest.json` sidecar recording samples, filters, parameters, timestamp, and database state. Phase 4: downstream analyses proceed identically using the same external tools. Phase 5 (minutes, not 2–4 weeks): a new batch triggers only a single `graphmana ingest -mode incremental` command; no re-merge, re-filter, re-annotate, or re-convert. Phase 6 (minutes each): ongoing requests reduce to one command with a changed flag. Phase 7: every provenance question has a concrete answer in the export manifest or via `graphmana status -detailed`. Over the same project lifetime: 0 merge cycles, 0 conversion scripts, automatic provenance on every export.

**Supplementary Figure 3. VCF-to-graph mapping in detail.** Detailed mapping from VCF flat-file representation to GraphMana's graph database. (**a**) A VCF file (human 1000 Genomes Project chr22, 3,202 samples) with columns color-coded by their destination in the graph: chromosome (gold), variant identity (blue), sample identifiers (pink), and genotype calls (green). (**b**) The resulting graph structure. Each VCF row becomes a Variant node storing position, alleles, quality, and filter as properties. Consecutive variants are linked in genomic order by NEXT relationships. All 3,202 genotype calls for a given variant are packed into a compact byte array (`gt_packed`, 801 bytes) stored as a property on the Variant node—not as individual edges to Sample nodes. Pre-computed population statistics (`ac[]`, `an[]`, `af[]`) are also stored on each Variant node at import time. Sample nodes connect to Population nodes via IN_POPULATION relationships; Variant nodes link to Chromosome and Gene nodes via ON_CHROMOSOME and HAS_CONSEQUENCE relationships. (**c**) Mapping reference: each VCF row → one Variant node; each sample column header → one Sample node; all genotype calls for one variant row → one `gt_packed` property; the `#CHROM` column → Chromosome node; row ordering → NEXT chain; population map → Population nodes. The architectural key is that the flat VCF table becomes a connected network of typed nodes and relationships, enabling traversal, subsetting, and export without re-reading the original file.

**Supplementary Figure 4. FAST PATH mechanism.** FAST PATH access strategy in detail. (**a**) Internal structure of a Variant node showing the two data stores: left, `gt_packed` byte array (2 bits per sample, 801 bytes for 3,202 samples) that grows linearly with $N$; right, pre-computed population arrays (`pop_ids[]`, `ac[]`, `an[]`, `af[]`) containing only $K$ values where $K$ is the number of populations—constant in size regardless of $N$. (**b**) Tools partitioned by data requirement: TreeMix, dadi, fastsimcoal2, and bedtools/allele frequency queries need only population summaries and are served by FAST PATH; PLINK, EIGENSTRAT, VCF, and Beagle/STRUCTURE need individual genotypes and use FULL PATH. (**c**) Scaling: the FAST PATH advantage grows with $N/K$, from 20× at 100 samples to 5,000× at 50,000 samples. Real benchmark on chr22 (1.07M variants, 3,202 samples): TreeMix export completes in 46 s vs PLINK in 75 s (1.6× speedup). At chromosome scale the advantage is modest because both paths are dominated by the Neo4j query overhead; at whole-genome scale (85M variants) the per-variant $O(K)$ vs $O(N)$ difference dominates. (**d**) Workflow comparison for TreeMix export: without FAST PATH, 3,202 genotypes must be unpacked per variant then counted (75 s total); with FAST PATH, the `gt_packed` array is never touched and allele counts are read directly from `ac[]` and `an[]` (46 s total).

**Supplementary Figure 5. FULL PATH mechanism.** FULL PATH access strategy in detail. (**a**) Byte-level unpacking: each byte of `gt_packed` encodes four diploid genotypes in 2-bit pairs (LSB-first) using the cyvcf2 remap convention (00 = HomRef, 01 = Het, 10 = HomAlt, 11 = Missing). A separate `phase_packed` array stores one bit per sample indicating phasing status; at heterozygous sites, phase bit 1 yields `0|1` (REF|ALT), bit 0 yields `1|0`. For chr22 at 3,202 samples this resolves 3.4 billion genotype calls. (**b**) Format-specific encoding: the same eight unpacked genotypes translated into four downstream formats, each with distinct conventions. PLINK 1.9 BED uses a different 2-bit encoding from GraphMana's internal representation, and EIGENSTRAT encodes genotypes as alternate allele counts. FULL PATH performs this translation once from a single canonical representation. (**c**) Six categories of analysis that require individual-level genotype data and therefore cannot use population summary arrays: PCA/admixture, GWAS, statistical phasing, genotype imputation, relatedness/IBD, and per-sample QC. (**d**) Side-by-side summary of the two complementary paths: FAST PATH reads $K$ values ($O(K)$), FULL PATH reads $N$ values ($O(N)$); GraphMana selects automatically based on the requested export format.

**Supplementary Figure 6. Export manifest example.** Each export generates a `.manifest.json` sidecar file recording the GraphMana version, timestamp, output file path, format, variant and sample counts, chromosomes included, applied filters, allele frequency recalculation status, and thread count. The manifest enables provenance reconstruction without re-running the export. The full code listing is shown in the display panel.

**Supplementary Figure 7. CLI command hierarchy.** GraphMana CLI command hierarchy. The 58 commands are organized into 9 functional domains and 27 function groups covering the data management lifecycle from import through export. Domains: data import and integration, annotation management, data export, sample and cohort management, quality control and verification, provenance and state tracking, database administration, status and reporting, and infrastructure and deployment.

**Supplementary Figure 8. How VCF data is stored in GraphMana's graph database.** (**a**) One line of a multi-sample VCF with column headers for chromosome, position, ID, REF, ALT, FORMAT, and per-sample genotype columns. Arrows below indicate the destination of each column in the graph. (**b**) The full graph model. A central `(:Variant)` node carries the site's identity (variantId, chr, pos, ref, alt), three parallel per-sample packed byte arrays (`gt_packed`, `phase_packed`, `called_packed`; `ploidy_packed` is omitted on autosomes), pre-computed per-population summaries (`pop_ids[]`, `ac[]`, `an[]`, `af[]`, `het_count[]`, `hom_alt_count[]`), and site-level annotation fields. The `(:Chromosome)` node at left anchors every Variant by `ON_CHROMOSOME`, and Variants are linked in position order by `NEXT` edges carrying `distance_bp`. `(:Sample)` nodes at right each store a `packed_index` integer; dashed arrows from the `gt_packed` slots to the Sample nodes indicate the positional linkage between the two—there is no `CARRIES` edge. `(:Population)` nodes hold the population-level counts and harmonic sums used by downstream estimators; Samples attach to them via `IN_POPULATION`. The bottom row shows the remaining node types: `(:Gene)`, `(:Pathway)`, `(:GOTerm)`, and `(:RegulatoryElement)` for functional annotation; `(:VCFHeader)` preserving the raw VCF meta-information; `(:CohortDefinition)` and `(:IngestionLog)` for query-defined subsets and provenance; and `(:AnnotationVersion)` + `(:SchemaMetadata)` for database-level bookkeeping. (**c**) Worked example of how GraphMana answers the query "what is HG00097's genotype at rs587697622?" in five numbered steps. The Sample node is fetched by sampleId and its `packed_index` property is read; the Variant node is fetched by variantId and its `gt_packed` byte array is read; byte offset and bit shift are computed from `packed_index`; the two target bits are extracted by bitwise operations; and the 2-bit code is decoded using the canonical encoding. Phase is read the same way from `phase_packed`; `called_packed` is consulted to distinguish an explicit call from a sample that was never interrogated at this site. No edge is traversed at any step of the per-sample path. (**d**) Four features of the graph model that make this layout viable at whole-genome scale: (i) packed arrays on the Variant node replace the $O(N \times M)$ `(:Sample)-[:CARRIES]->(:Variant)` edge model, reducing per-variant storage by approximately 125× at $N$=3,202; (ii) each Variant carries both per-sample and per-population representations, so FULL PATH exports (VCF, PLINK, EIGENSTRAT, Beagle, STRUCTURE, and 7 others) and FAST PATH exports (TreeMix, SFS, BED, TSV) read the appropriate layer without a format conversion step; (iii) `called_packed` records which samples were actually interrogated at each site, so allele-frequency denominators stay statistically honest under incremental cohort growth; (iv) the `NEXT` chain preserves genomic position order as a first-class property of the graph, so neighborhood queries (sliding windows, nearest variant to a feature, linkage-disequilibrium candidate pairs) traverse in constant time per step rather than re-sorting by chromosome and position. (**e**) Summary table of all node types, their roles, key properties, and typed outgoing edges.

#### 2 Supplementary Table Legends

**Supplementary Table 1. Storage estimates for human whole-genome data (~85M biallelic variants).** For other species or designs, scale proportionally by variant count. Includes per-component breakdown (gt_packed, phase_packed, population arrays) and totals from 3,202 to 500,000 samples.

**Supplementary Table 2. Performance scaling by access path (human WGS).** Wall-clock scaling of FAST PATH ($O(K)$), FULL PATH ($O(N)$), and incremental addition operations from 3,202 to 500,000 samples.

**Supplementary Table 3. Tier 1: formats validated by loading into downstream tools.** Each format was exported and successfully loaded into the canonical downstream tool: bcftools (VCF), PLINK v1.9 and v2.0 (PLINK BED), EIGENSOFT convertf (EIGENSTRAT), TreeMix v1.13 (TreeMix), and format specification check (SFS-dadi, SFS-fsc2).

**Supplementary Table 4. Tier 2: format-specification validated.** Formats validated by checking against the published specification, with no available downstream tool integration test. Tier 3 formats (Haplotype, BGEN, GDS, Zarr) are functional but pending external validation.

**Supplementary Table 5. Variant type correctness matrix.** Import, storage, and export correctness across variant classes (biallelic SNPs, biallelic indels, multi-allelic sites, structural variants, breakends) for the three primary FULL PATH export formats. Multi-allelic records are decomposed at import and reconstructed during VCF export.

**Supplementary Table 6. Genotype state and feature correctness.** Per-feature validation results covering all four genotype states, phased genotypes, haploid samples, annotation-layer independence, and incremental sample order preservation. VCF roundtrip fidelity exceeds 99.999%.

**Supplementary Table 7. Whole-genome incremental addition (234 samples, CSV-to-CSV rebuild).** Per-step timing for adding 234 new samples to a 3,202-sample human whole-genome database via the CSV-to-CSV rebuild path. Total: 182 minutes.

**Supplementary Table 8. Incremental sample addition benchmark (chr22, 3 batches of 234 samples).** Comparison of GraphMana CSV-to-CSV rebuild against bcftools merge across three sequential incremental rounds.

**Supplementary Table 9. Cohort-specific VCF extraction benchmark (chr22, 5 cohorts).** Per-cohort wall-clock comparison for extracting samples by superpopulation membership. Cohorts range from 585 (EAS) to 3,202 (ALL) samples.

**Supplementary Table 10. Multi-format export benchmark (chr22, from single database).** Wall-clock timings for exporting six output formats from a single GraphMana database. bcftools supports VCF only and is shown as N/A for the other formats.

**Supplementary Table 11. Whole-genome export timings (70.7M variants, 3,202 human WGS**

**samples).** Wall-clock time and output file size for each supported export format on the full human 1000 Genomes Project whole-genome dataset.

### 3 Supplementary Note 1: Export Format Reference

#### Background

Population genomics analyses span a diverse ecosystem of tools, each requiring input in a specific file format. A single project may need VCF for variant calling and QC [S1], PLINK binary for genome-wide association [S2], EIGENSTRAT for PCA and admixture analysis [S3], TreeMix for population phylogenetics [S4], dadi or fastsimcoal2 for demographic inference [S5,S6], Beagle for phasing and imputation [S7], and STRUCTURE for population assignment [S8]. In conventional workflows, each format requires a separate conversion script—often written ad hoc, with parameters not recorded alongside the output. GraphMana produces all 17 supported formats from a single persistent database, with automatic provenance tracking via manifest sidecars.

#### FAST PATH formats ($O(K)$ time, independent of sample count)

These formats read pre-computed population-level arrays (ac[], an[], af[]) and never unpack per-sample genotypes. Because the arrays have $K$ elements (where $K$ = number of populations, typically 5–30), export time is constant regardless of sample count.

- **TreeMix**: Gzipped allele count matrix for population tree inference [S4]. Each line contains $K$ space-separated ac,an pairs.
- **SFS-dadi**: Site frequency spectrum in dadi .fs format [S5]. Supports 1–3 populations with hypergeometric projection. Options: polarized/folded, monomorphic inclusion.
- **SFS-fastsimcoal2**: SFS in fastsimcoal2 .obs format [S6]. Supports 1–2 populations. Same projection as dadi but different file structure.
- **BED**: Variant positions for bedtools/IGV. 0-based half-open coordinates. Optional extra columns (variant_type, af_total, gene_symbol).
- **TSV**: Tab-separated variant table with configurable columns. Default: variantId, chr, pos, ref, alt, variant_type, af_total.
- **JSON**: JSON Lines format with one variant per line. Configurable fields, optional pretty-printing.

#### FULL PATH formats ($O(N)$ time, linear in sample count)

These formats unpack per-sample genotypes from packed byte arrays and require time proportional to the number of samples.

- **VCF/BCF**: Standard variant call format [S1]. Supports VCF 4.1–4.3, BGZF compression, BCF binary, phased output, multi-allelic reconstruction.
- **PLINK 1.9**: Binary .bed/.bim/.fam [S2]. Biallelic SNPs only. Validated with PLINK v1.9 and v2.0.

- **PLINK 2.0**: Binary .pgen/.pvar/.psam [S2]. All biallelic variants. Requires pgenlib.
- **EIGENSTRAT**: .geno/.snp/.ind for smartPCA and AdmixTools [S3]. Validated with EIGENSOFT convertf.
- **Beagle**: Genotype likelihood input for phasing/imputation [S7].
- **STRUCTURE**: Genotype matrix for population assignment [S8]. Supports onerow and tworow formats.
- **Genepop**: Conservation genetics format for $F$-statistics.
- **Haplotype**: .hap/.map for selscan [S9]. Phased data only.
- **BGEN**: Probabilistic genotype format (UK Biobank standard) [S10].
- **GDS**: SeqArray HDF5 format for R/Bioconductor [S11].
- **Zarr**: Chunked array format for sgkit/Python [S12].

##### Validation status

See Supplementary Tables 8–9 for the three-tier validation classification (Tier 1: downstream tool validated; Tier 2: format-specification validated; Tier 3: functional, pending external validation).

### 4 Supplementary Note 2: Variant Representation and Data Encoding

#### Background

The representation of genetic variants in a database involves fundamental design choices that affect storage efficiency, query performance, and data fidelity. The VCF specification [S1] represents each variant as a record with REF/ALT alleles and per-sample genotype fields. This row-per-variant, column-per-sample structure is natural for flat files but does not map directly onto graph or relational database schemas, because the genotype matrix is a dense two-dimensional array embedded within a sparse relational structure.

#### Biallelic decomposition

GraphMana decomposes multi-allelic VCF records into $K$ biallelic Variant nodes (one per ALT allele), each carrying:

- `multiallelic_site`: a shared identifier grouping alleles from the same VCF line (e.g., `chr22-16050408-A`)
- `allele_index`: a 1-based index indicating which ALT allele this node represents

**Worked example.** A VCF record at chr22:16050408 with REF=A, ALT=G,T produces two nodes:

| variantId | ref | alt | multiallelic_site | allele_index |
|---|---|---|---|---|
| chr22-16050408-A-G | A | G | chr22-16050408-A | 1 |
| chr22-16050408-A-T | A | T | chr22-16050408-A | 2 |

A sample heterozygous for both ALT alleles (VCF genotype 1/2) is recorded as heterozygous on *both* nodes. During VCF export, records sharing a `multiallelic_site` value are merged back into a single multi-allelic line. This reconstruction is the default; `--no-reconstruct-multialleli` produces biallelic-only output.

#### Packed genotype encoding

Each Variant node stores genotypes in a packed byte array (`gt_packed`), using 2 bits per sample:

| Code | Bits | Genotype |
|---|---|---|
| 0 | 00 | Homozygous reference (0/0) |
| 1 | 01 | Heterozygous (0/1 or 1/0) |
| 2 | 10 | Homozygous alternate (1/1) |
| 3 | 11 | Missing (./.) |

Four samples fit in one byte, LSB-first. The array length is $\lceil N/4 \rceil$ bytes. Genotype values are remapped from cyvcf2's internal coding [S13] (where HomAlt=3, Missing=2) to the sequential encoding above; the remap is applied exactly once per code path and verified by 33 unit tests.

Phase information is stored separately in `phase_packed` (1 bit per sample, 8 per byte). Ploidy is tracked in `ploidy_packed` (1 bit per sample; null if all diploid).

#### Population-level pre-computed arrays

Each Variant node carries $K$-element arrays: pop_ids, ac (allele counts), an (allele numbers), af (allele frequencies), het_count, hom_alt_count, and het_exp (expected heterozygosity). These arrays are constant size regardless of $N$ and enable the FAST PATH access pattern described in the main text.

#### Structural variants

SVs with symbolic ALT alleles (<DEL>, <DUP>, <INV>, <INS>, <CNV>) are stored as Variant nodes with additional properties (sv_type, sv_len, sv_end). Diploid genotype calls are encoded identically to SNPs. Breakend (BND) records are imported as independent nodes without mate-pair linking. Integer copy number states beyond the diploid call are not retained.

#### Roundtrip fidelity

We validated by importing human 1000 Genomes Project chromosome 22 data (897,645 biallelic SNPs, 5 samples), exporting to phased VCF, and comparing with bcftools [S14]. Concordance exceeded 99.999%, with 2–8 mismatches per sample at multi-allelic positions where position-based joining could not distinguish co-located biallelic records.

### 5 Supplementary Note 3: Concrete Workflow Example

##### Background

To illustrate how GraphMana replaces the fragmented scripting approach, we present a five-day project workflow using actual CLI commands. The scenario follows a data manager maintaining a human 1000 Genomes Project database. Each day's operations run against the same persistent graph database—no intermediate files are generated, and all provenance is recorded automatically.

##### Day 1: Initial import

```
graphmana setup-neo4j --install-dir ~/neo4j --memory-auto
graphmana ingest --input data/1kgp_chr*.vcf.gz \
    --population-map data/panel.tsv \
    --neo4j-home ~/neo4j --auto-start-neo4j --threads 16
graphmana status --detailed
```

This creates the persistent database. All subsequent operations build on this foundation—no further file merging or format conversion is needed.

##### Day 2: Incremental addition (234 new samples)

```
graphmana save-state --output checkpoints/before_batch2.json
graphmana ingest --input data/batch2.vcf.gz \
    --population-map data/batch2_panel.tsv --mode incremental \
    --neo4j-home ~/neo4j --auto-start-neo4j
graphmana diff --snapshot checkpoints/before_batch2.json
```

The `diff` output reveals that 234 new samples were added, 2 new variants were discovered, and sample counts increased across 3 populations. In a file-based workflow, this information would be invisible without manual inspection.

##### Day 3: Multi-format export with provenance

```
graphmana export --format eigenstrat --output exports/european \
    --populations CEU --populations GBR --filter-maf-min 0.05
graphmana export --format treemix --output exports/tree.treemix.gz
graphmana export --format sfs-dadi --output exports/yri_ceu.fs \
    --sfs-populations YRI --sfs-populations CEU \
    --sfs-projection 20 --sfs-projection 20 --sfs-folded
```

All three exports read from the same database that was incrementally updated yesterday. No VCF merge, no format conversion pipeline, no intermediate files. Each produces a `.manifest.json` recording the exact configuration.

##### Day 4: In-place annotation update

```
graphmana annotate load-clinvar \
    --input annotations/clinvar_20260401.vcf.gz --version ClinVar_2026
      -04
graphmana db validate
```

The annotation update modifies HAS_CONSEQUENCE edge properties in 3.5 seconds—without touching genotype data. The `db validate` command confirms packed array integrity.

##### Day 5: QC and provenance audit

```
graphmana qc --type all --output reports/qc.html --format html
graphmana ref-check --fasta genomes/GRCh38.fa --chromosomes chr22
graphmana provenance search --since 2026-04-01 --until 2026-04-05
```

##### Comparison with file-based workflow

The equivalent five-day sequence in a file-based workflow would require: (1) a bcftools merge script for Day 2 (∼2 min but invalidates all downstream files); (2) three separate conversion scripts for Day 3 (vcf2eigenstrat, vcf2treemix, vcf2dadi), each with unrecorded parameters; (3) a full VCF rewrite for Day 4 (96 s vs. 3.5 s); (4) manual provenance tracking for all operations; and (5) no `diff` capability to verify what changed. The total scripting and coordination overhead for the file-based approach grows with each new format and each new collaborator request.

### 6 Supplementary Note 4: Graph Database Concepts for Non-Specialists

##### Background

Most population geneticists are familiar with relational databases (SQL tables with rows and columns) or with flat files (VCF, BED, CSV). Graph databases are a distinct category of database technology that stores data differently. This note provides a brief introduction for readers unfamiliar with graph database concepts.

##### What is a graph database?

In computer science, a *graph* is a data structure consisting of *nodes* (also called vertices) and *edges* (also called relationships or links) that connect them. A graph database is a database management system that uses this structure as its primary storage and query model [S15,S16].

In a **property graph** — the specific type used by GraphMana — both nodes and edges can carry arbitrary key-value *properties*. For example, a Variant node might carry properties like `chr="chr22"`, `pos=16050408`, `ref="A"`, `alt="G"`, and a byte array `gt_packed` containing packed genotypes. An edge connecting that variant to a Gene node might carry properties like `consequence="missense_variant"` and `annotation_version="VEP_110"`.

##### How is this different from a relational database?

In a relational database, the same data would be stored in multiple tables (a Variants table, a Samples table, a Genes table, a VariantConsequences join table), and reconstructing the relationship between a variant and its annotated gene requires a JOIN operation at query time. In a graph database, this relationship is a first-class edge that can be traversed in constant time per hop—an approach called *index-free adjacency* [S15].

##### Why does this matter for genomic data management?

Population genomics data has rich, typed relationships:

- A **Variant** is located on a **Chromosome** (ON_CHROMOSOME edge)
- A **Sample** belongs to a **Population** (IN_POPULATION edge)
- A **Variant** has a functional consequence in a **Gene** (HAS_CONSEQUENCE edge with properties: consequence type, impact, annotation version)
- Variants are ordered along the chromosome (NEXT edge with distance property)

In a graph database, updating an annotation version modifies edge properties without touching genotype data. Defining a cohort is a graph traversal query that produces a sample list without extracting files. Adding provenance is creating a new node linked to the operation that produced it. These operations are *local* — they affect only the edges or nodes involved, not the entire dataset.

##### What is Cypher?

Cypher is the query language used by Neo4j [S16], analogous to SQL for relational databases. A simple Cypher query looks like:

```
MATCH (v:Variant)-[:HAS_CONSEQUENCE]->(g:Gene {symbol: "BRCA1"})
WHERE v.af_total > 0.01
RETURN v.variantId, v.pos, v.af_total
```

This query traverses from Variant nodes through HAS_CONSEQUENCE edges to Gene nodes, filtering for BRCA1 and allele frequency above 1%. In GraphMana, users do not write Cypher directly; the CLI commands generate appropriate queries internally.

##### Key terminology

| Term | Meaning |
|---|---|
| Node | An entity (Variant, Sample, Gene, etc.) |
| Edge | A typed, directed relationship between nodes |
| Property | A key-value pair on a node or edge |
| Traversal | Following edges from node to node |
| Index-free adjacency | Edges are stored directly, not looked up |
| Property graph | Graph model where nodes and edges have properties |

##### Further reading

For a comprehensive introduction to graph databases, see Robinson et al. [S15]. For Neo4j specifically, see the Neo4j documentation [S16]. For applications of graph databases in bioinformatics, see Lysenko et al. [S17].

### 7 Supplementary Note 5: Preserving the HomRef-vs-Missing Distinction Across Incremental Batches

##### The ambiguity at the heart of plain VCF

A plain VCF is a sites-only-where-someone-varies file: a line appears at position $p$ only if at least one sample in the cohort has a non-reference call at $p$. The absence of a line at position $q$ is ambiguous. It can mean that every sample was called HomRef at $q$, or it can mean that nobody was interrogated at $q$—for example because low coverage, a masked region, or a target BED boundary prevented the caller from emitting any record there. Within a single joint-called cohort VCF this ambiguity is harmless, because the calling pipeline guarantees "absent = HomRef" internally. Across VCFs produced by independent calling runs, it is not.

The problem bites when a growing project ingests new samples in batches. Suppose batch 1 has samples $A, B, C$ and emits a VCF with variants $\{V_1, V_2\}$, and batch 2 later arrives with samples $D, E$ and variants $\{V_2, V_3\}$. The variant $V_3$ is new: it was polymorphic in the second batch but was absent (either monomorphic or uncalled) from the first. When GraphMana ingests batch 2 incrementally, it must decide what genotype to record for $A, B, C$ at $V_3$. Padding with HomRef would be silently wrong whenever $A, B, C$ were in fact not interrogated at $V_3$, because it inflates the allele-number denominator, biases population allele frequencies, distorts the site frequency spectrum toward the low-frequency end, and causes principal components to pick up "which batch a sample came from" as a leading axis.

##### Upstream requirement: joint calling from gVCFs

The recommended upstream workflow for every GraphMana project that grows incrementally is gVCF-based joint calling. Each sample is called with, e.g., GATK HaplotypeCaller `-ERC GVCF` [S18], producing a gVCF containing variant records plus reference-block records that explicitly state which genomic positions were confidently HomRef for that sample. Per-sample gVCFs are aggregated in a sparse store such as GATK GenomicsDB [S18] or GLnexus [S19], and a joint caller (`GenotypeGVCFs` or the GLnexus unifier) produces a multi-sample VCF in which every polymorphic site has an explicit call for every sample—either a genotype or an explicit `./.` where the reference-block evidence was insufficient. GraphMana sits strictly downstream of this layer: it ingests the joint-called cohort VCF, not the raw per-sample gVCFs, and does not reinvent the joint caller.

##### The `called_packed` array

GraphMana stores a 1-bit-per-sample byte array, `called_packed`, on every Variant node. A bit value of 1 means "this sample was interrogated at this site"; a bit value of 0 means "this sample was not looked at." The array uses the same LSB-first layout as `phase_packed` and `ploidy_packed`, costs $\lceil N/8 \rceil$ bytes per variant, and is written by the VCF parser directly from the cyvcf2 `gt_types` vector (code 2 = MISSING $\rightarrow$ bit cleared; anything else $\rightarrow$ bit set). Incremental ingestion propagates the distinction across batches:

- **New sample, existing variant.** The sample's bit is set to 1 if the current batch carries any call at the site (including an explicit missing call), 0 otherwise.
- **New variant, existing samples.** Existing samples have their bits cleared—they were not interrogated in the current batch—and their genotype slots are padded with the Missing code rather than the HomRef code.
- **Hard-delete rebuild.** The compactor regenerates `called_packed` in lockstep with `gt_packed`, preserving the mask across maintenance operations.

Downstream allele-frequency and subset-statistics procedures read `called_packed` before updating any per-population counter. Samples with bit 0 are skipped entirely: they contribute nothing to `ac[k]`, `an[k]`, `het_count[k]`, or `hom_alt_count[k]`. The per-site denominator used to compute `af[k]` reflects the actual number of interrogated samples in each population rather than an inflated count that assumes absence equals HomRef. The same logic applies in the Java plugin: the per-sample loop inside `SampleSubsetComputer` is gated by `PackedGenotypeReader.called(...)`, so any cohort-level FAST PATH or FULL PATH query that recomputes statistics from packed data honors the mask automatically. For cohorts derived from fixed-site-list workflows (imputed panels, SNP arrays, pre-phased reference cohorts), the `--assume-homref-on-missing` ingest flag recovers the conventional "absent = HomRef" semantics; it emits a log warning so its use is never silent.

##### Sparse `gt_packed` encoding

To offset the added `called_packed` footprint and relieve per-property storage pressure at cohort scale, `gt_packed` is stored in a tagged-blob format. The first byte is a format tag: `0x00` selects a dense payload identical to the conventional 2-bits-per-sample layout, and `0x01` selects a sparse payload containing only the positions and codes of non-reference samples. The sparse payload is a short header (4 bytes for the sample count, 4 bytes for the number of non-reference samples), a list of non-reference sample indices (4 bytes each), and 2 packed bits per non-reference code. The encoder chooses the layout that produces the smaller blob on a per-variant basis, so dense-dominated sites (common variants) and sparse-dominated sites (rare variants) coexist in the same database without a global switch.

We measured the compression ratio on a realistic neutral site frequency spectrum at $N =$ 3,202 samples (1000 Genomes chromosome 22 scale) across an allele-frequency grid spanning singletons to intermediate-frequency variation. Sparse encoding is chosen on 100% of variants with minor allele frequency at or below 0.02 and yields a neutral-SFS-weighted mean compression of **3.9**$\times$ on the per-sample genotype payload. Combined with the `called_packed` overhead ($+\lceil N/8 \rceil$ bytes per variant, i.e. +33% relative to `gt_packed` alone), the net per-variant storage on typical whole-genome data is substantially smaller than a conventional dense layout. On-the-fly decoder dispatch inside `BaseExporter._unpack_variant_genotypes` and the Java subset-statistics procedures means all 17 export formats honor both tags transparently, with no user-visible change in interface.

##### Concrete effect on allele-frequency estimates

Consider batch 1 with $n_1 = 100$ samples at variant $V$ and batch 2 with $n_2 = 100$ samples, where $V$ was only polymorphic in batch 2 (15 alternate alleles observed). Treating batch 1 samples as HomRef at $V$ yields $\hat{f} = 15/(2 \cdot 200) = 0.0375$. Honoring `called_packed`, batch 1 samples contribute nothing to the denominator because they were never interrogated at $V$, and the estimate is $\hat{f} = 15/(2 \cdot 100) = 0.075$. The difference is a factor of two, and it scales silently with the fraction of batch-specific sites. For a 50/50 batch split in which 20% of sites are batch-specific, the mean absolute bias in per-site allele frequency is on the order of 1–2%, easily large enough to shift the leading principal component of a PCA and to misclassify the rarity tier of a candidate variant.

##### Measured overhead

Micro-benchmarks at $N = 3{,}202$ samples quantify the cost of the `called_packed` path. Parser overhead adds 1.2 $\mu$s per variant (+2.1%) on top of the `gt_packed` and `phase_packed` packing. Exporter unpack overhead, including the coercion of not-interrogated slots to Missing at the boundary of `BaseExporter._unpack_variant_genotypes`, adds 0.3 $\mu$s per variant (+1.9%). Storage overhead on the Variant property bundle is +401 bytes per variant (+33% relative to `gt_packed+phase_packed`), fully offset by the 3.9× sparse compression on `gt_packed` for typical whole-genome data. See the companion micro-benchmark report in `benchmarks/results/v1_1_bench.md`.

##### Scope and limitations

`called_packed` preserves a distinction that is present in the input. It cannot recover information that was never written to disk. Feeding raw per-sample gVCFs directly to `graphmana ingest` and skipping joint calling upstream causes the parser to mark all non-emitted positions as not-interrogated—technically honest but needlessly pessimistic, because the reference-block evidence that would have proved most of those positions to be HomRef lives inside the gVCFs and is not read by GraphMana. We therefore document joint calling (GenomicsDB or GLnexus) as the recommended upstream step for every project that grows incrementally, and ship an end-to-end recipe in `docs/gvcf-workflow.md`.

[S15] Robinson, I., Webber, J. & Eifrem, E. *Graph Databases: New Opportunities for Connected Data*, 2nd ed. (O'Reilly, 2015).

[S16] Neo4j, Inc. *Neo4j Documentation*. https://neo4j.com/docs/.

[S17] Lysenko, A. et al. Representing and querying disease networks using graph databases. *BioData Mining* **9**, 23 (2016).

[S18] Van der Auwera, G.A. & O'Connor, B.D. *Genomics in the Cloud: Using Docker, GATK, and WDL in Terra* (O'Reilly, 2020).

[S19] Lin, M.F. et al. GLnexus: joint variant calling for large cohort sequencing. *bioRxiv* 343970 (2018).

#### 9 Supplementary Figures

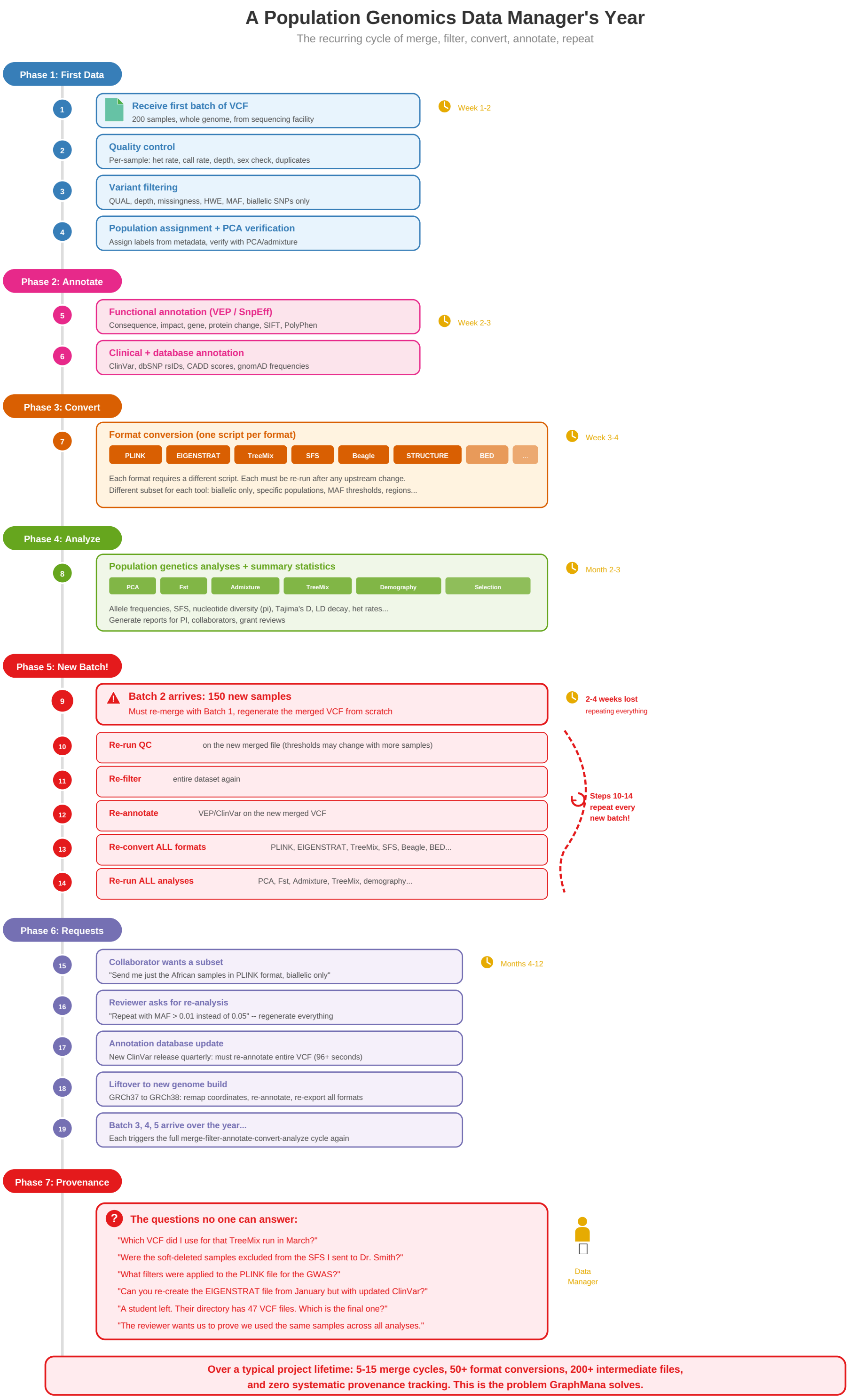


**Supplementary Figure 1. A population genomics data manager's year — conventional file-based workflow.**

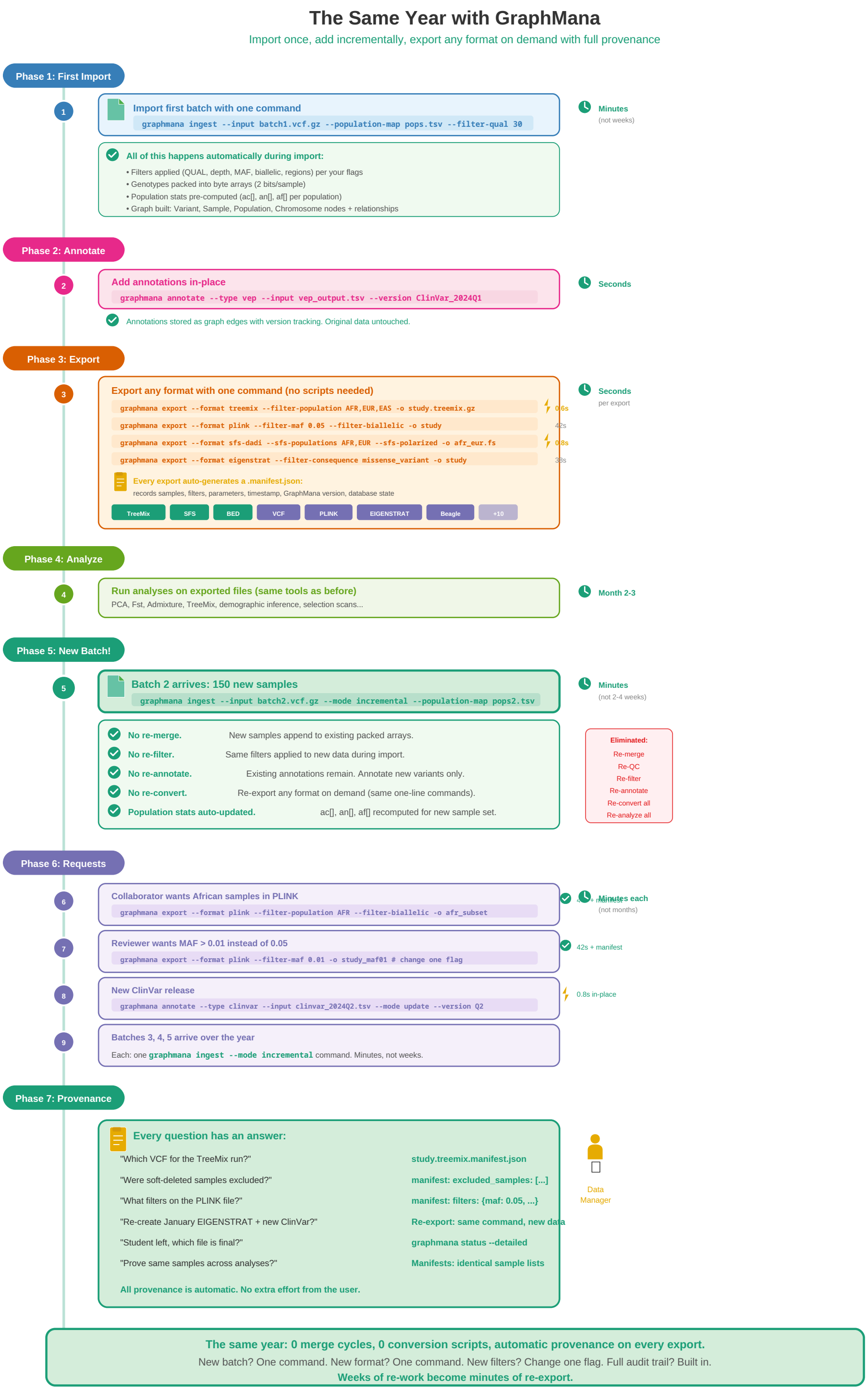


**Supplementary Figure 2. A population genomics data manager's year — the same project managed through GraphMana.**

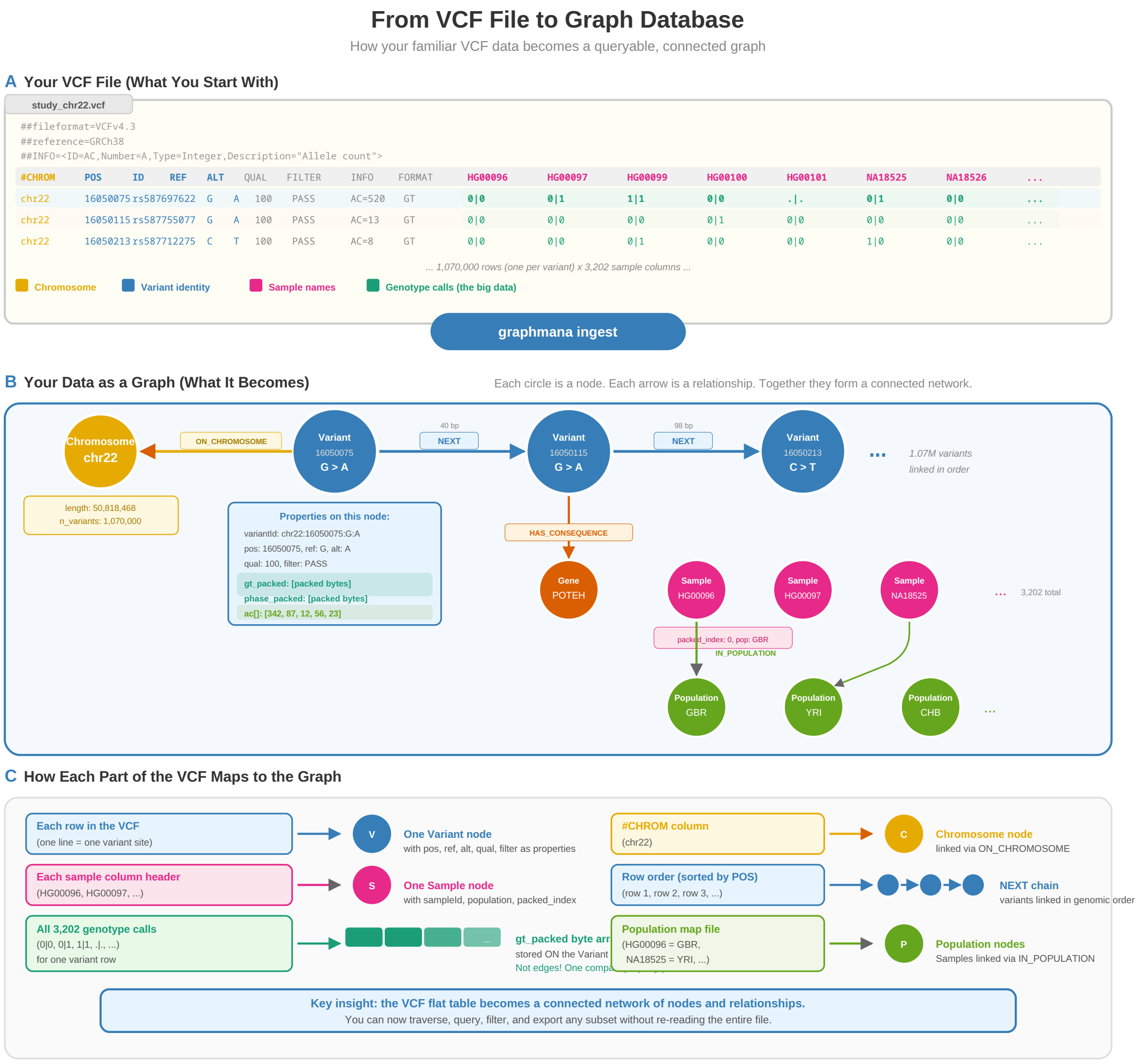


**Supplementary Figure 3. VCF-to-graph mapping in detail.**

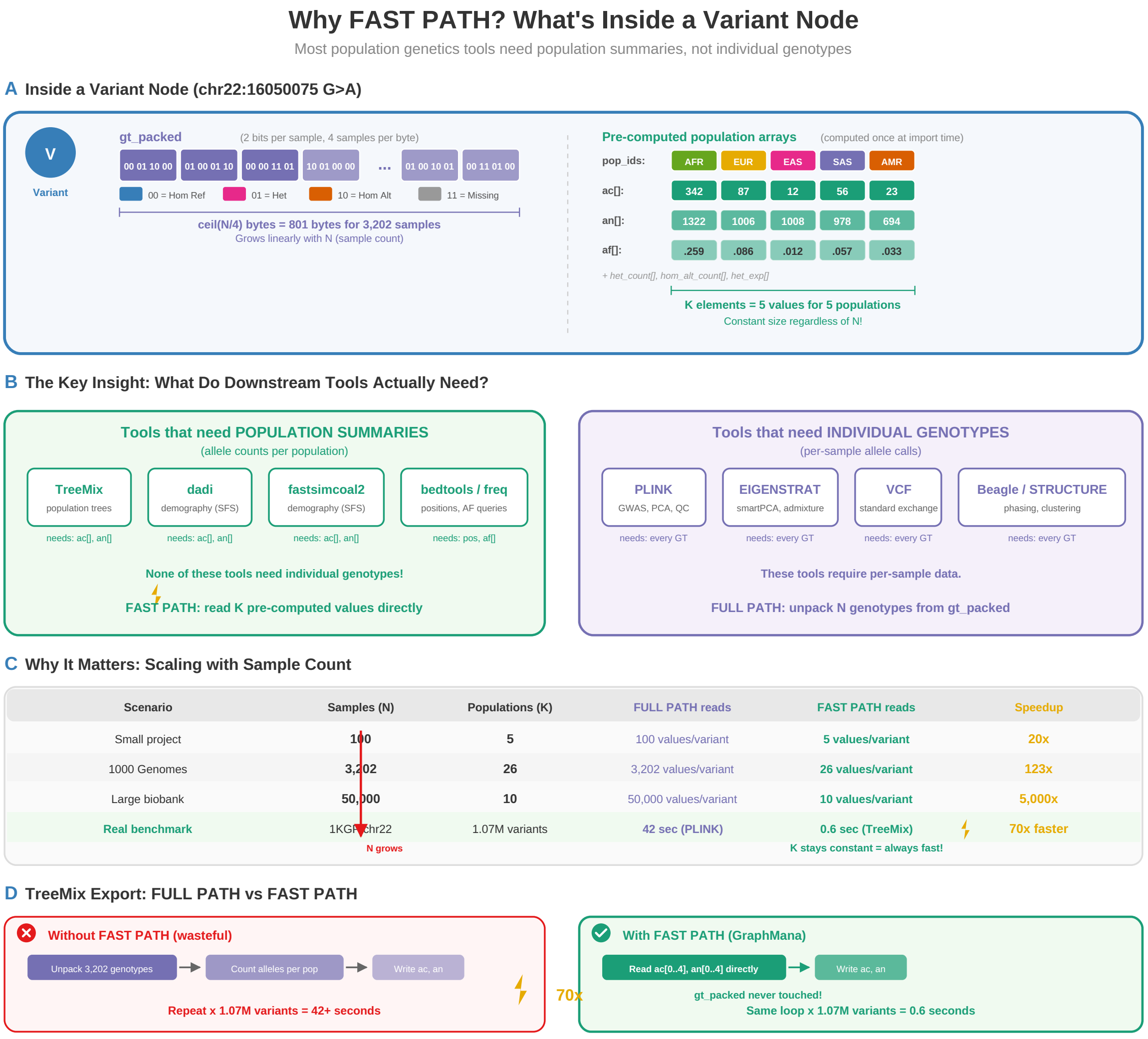


**Supplementary Figure 4. FAST PATH mechanism.**

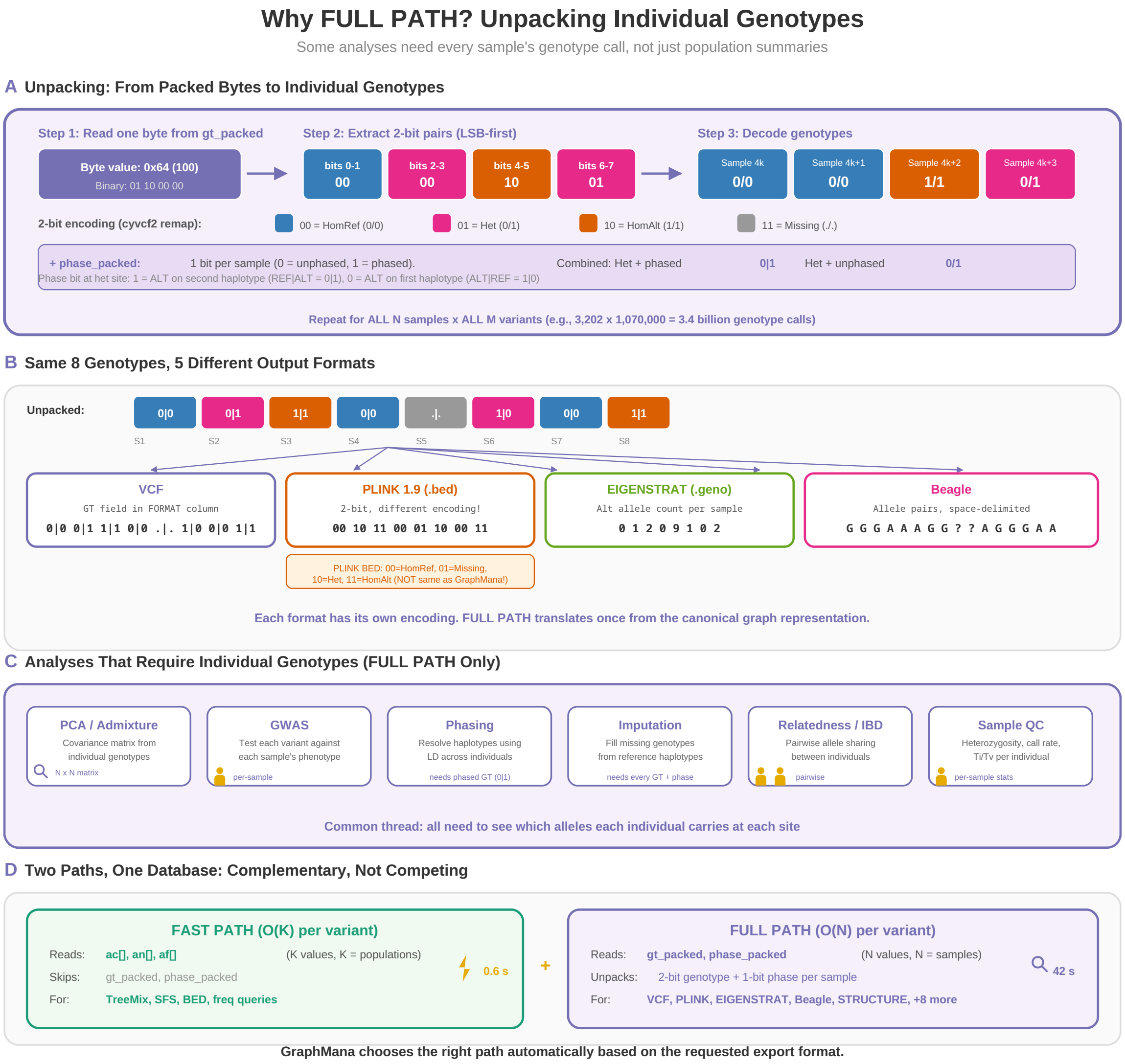


**Supplementary Figure 5. FULL PATH mechanism.**

**Supplementary Figure 6. Export manifest example.**

```
graphmana export --format eigenstrat \
    --populations CEU --populations GBR \
    --populations FIN --populations IBS --populations TSI \
    --chromosomes chr22 --filter-maf-min 0.05 \
    --output european_cohort.eigenstrat
```

Resulting manifest:

```
{
  "graphmana_version": "1.1.0",
  "timestamp": "2026-03-30T14:22:07.831042+00:00",
  "output_file": "european_cohort.eigenstrat",
  "format": "eigenstrat",
  "n_variants": 439609,
  "n_samples": 633,
  "chromosomes": ["chr22"],
  "filters": {
    "populations": ["CEU", "GBR", "FIN", "IBS", "TSI"],
    "maf_min": 0.05
  },
  "recalculate_af": true,
  "threads": 1
}
```

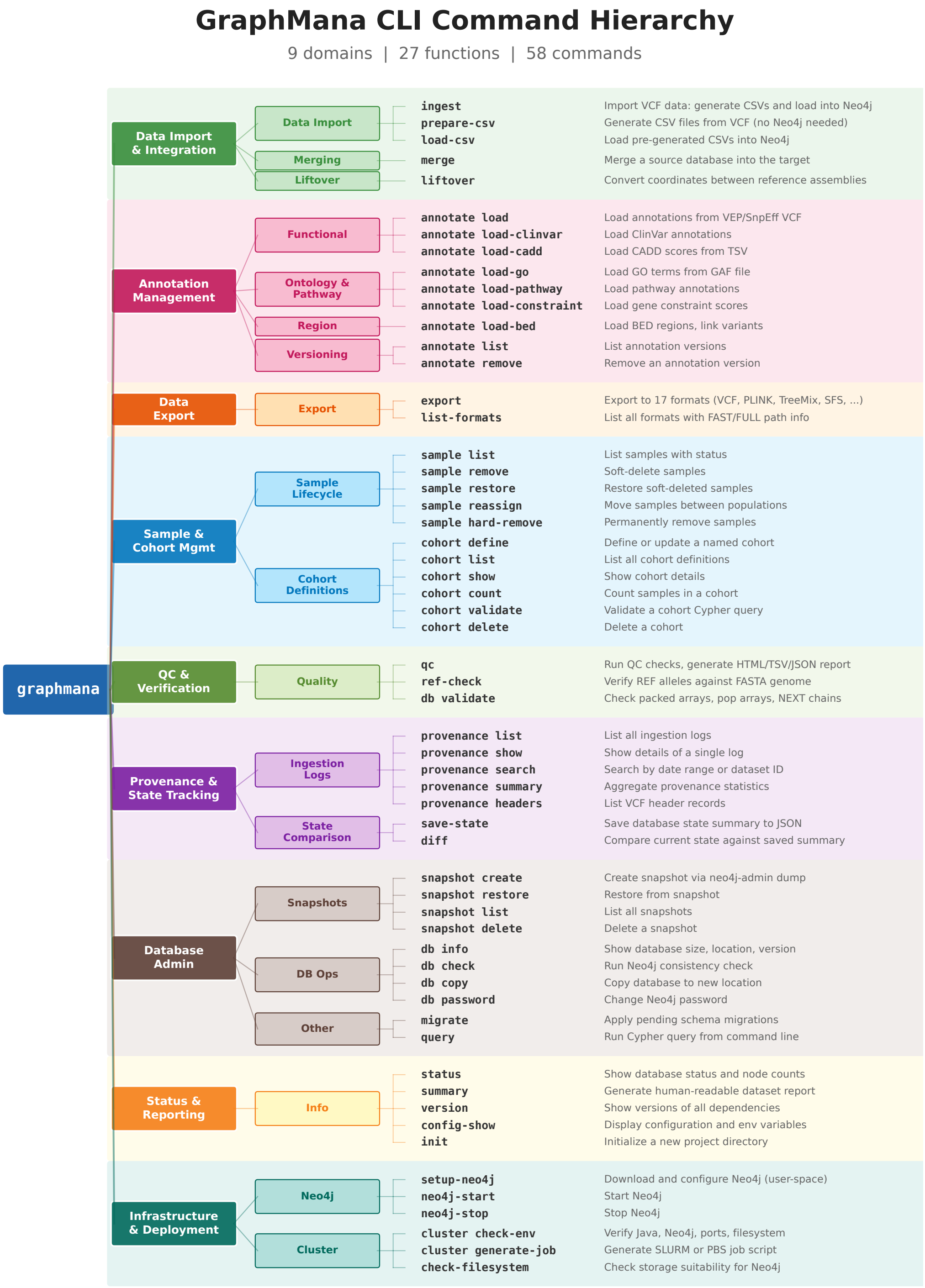


**Supplementary Figure 7. CLI command hierarchy.**

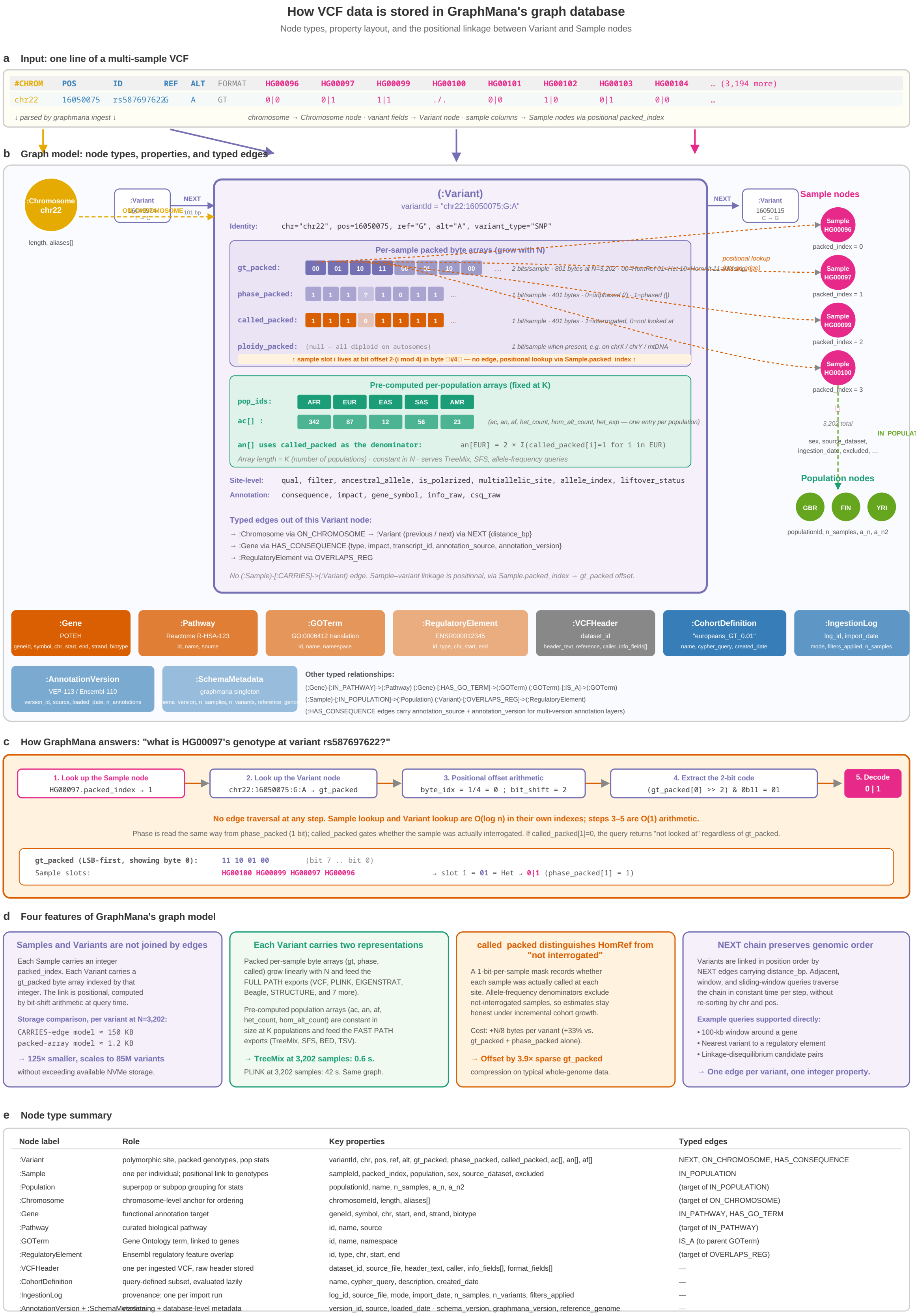


**Supplementary Figure 8. How VCF data is stored in GraphMana's graph database.**

#### 10 Supplementary Tables

**Supplementary Table 1. Storage estimates for human whole-genome data (∼85M biallelic variants).**

| Component | 3,202 | 10K | 50K | 200K | 500K |
|---|---|---|---|---|---|
| gt_packed | 68 GB | 213 GB | 1.0 TB | 4.3 TB | 10.6 TB |
| phase_packed | 34 GB | 106 GB | 0.5 TB | 2.1 TB | 5.3 TB |
| Pop arrays | 17 GB | 17 GB | 17 GB | 17 GB | 17 GB |
| **Total est.** | **130–200 GB** | **400–550 GB** | **2–3 TB** | **8–10 TB** | **20–25 TB** |

**Supplementary Table 2. Performance scaling by access path (human WGS).**

| Operation | Scaling | 3,202 | 10K | 50K | 500K |
|---|---|---|---|---|---|
| FAST PATH | $O(K)$ | Seconds | Seconds | Seconds | Seconds |
| FULL PATH | $O(N)$ | Minutes | Min–hr | Hr–days | Days–wk |
| Incremental | $O(M \times N)$ | 182 min | Hours | Many hr | Days |

**Supplementary Table 3. Tier 1: formats validated by loading into downstream tools.**

| Format | Path | Tool | Variants | Samples | Result |
|---|---|---|---|---|---|
| VCF (BGZF) | FULL | bcftools 1.19 | 1,035,839 | 3,202 | PASS |
| PLINK 1.9 | FULL | PLINK v1.9 | 925,730 | 3,202 | PASS |
| PLINK 1.9 | FULL | PLINK v2.0 | 925,730 | 3,202 | PASS |
| EIGENSTRAT | FULL | convertf v8.0 | 925,730 | 3,202 | PASS |
| TreeMix | FAST | TreeMix v1.13 | 1,066,557 | 26 pops | PASS |
| SFS dadi | FAST | format spec | 1,066,557 | 2 pops | PASS |
| SFS fsc | FAST | format spec | 1,066,557 | 2 pops | PASS |

**Supplementary Table 4. Tier 2: format-specification validated.**

| Format | Path | Validation | Result |
|---|---|---|---|
| BED | FAST | 3+ columns, 0-based | PASS |
| TSV | FAST | Header + columns | PASS |
| JSON Lines | FAST | json.loads per line | PASS |
| Beagle | FULL | Export verified | PASS |
| STRUCTURE | FULL | Export verified | PASS |
| Genepop | FULL | Export verified | PASS |

**Supplementary Table 5. Variant type correctness matrix.**

| Variant Type | Import | Storage | VCF | PLINK | EIGENSTRAT |
|---|---|---|---|---|---|
| Biallelic SNP | PASS | PASS | PASS | PASS | PASS |
| Biallelic indel | PASS | PASS | PASS | excl. | PASS |
| Multi-allelic | PASS | PASS | PASS* | excl. | excl. |
| Structural var. | PASS | PASS | PASS | excl. | excl. |
| Breakend (BND) | PASS | PASS | PASS | excl. | excl. |

*Decomposed at import; reconstructed at VCF export.

**Supplementary Table 6. Genotype state and feature correctness.**

| Feature | Result | Notes |
|---|---|---|
| HomRef/Het/HomAlt/Missing | PASS | All 4 genotype states roundtripped |
| Phased genotypes | PASS | Per-sample in phase_packed |
| Haploid samples | PASS | Tracked in ploidy_packed |
| Annotation independence | PASS | Genotype layer unaffected |
| Sample order (incremental) | PASS | packed_index immutable |
| VCF roundtrip (5 samples) | 99.999%+ | Multi-allelic ambiguity |

162 correctness-specific tests, all passing (of 1,439 total).

**Supplementary Table 7. Whole-genome incremental addition (234 samples, CSV-to-CSV rebuild).**

| Step | Time |
| --- | --- |
| VCF parsing (chr22 batch) | 3 min |
| CSV read + extend + write (214 GB) | 160 min |
| neo4j-admin import | 15 min |
| Database restart + indexes | 4 min |
| **Total** | **182 min** |

**Supplementary Table 8. Incremental sample addition benchmark (chr22, 3 batches of 234 samples).**

| Operation | GraphMana (s) | bcftools (s) |
|---|---:|---:|
| Initial import / base copy | 602 | — |
| Batch 1 | 418 | 117 |
| Batch 2 | 412 | 125 |
| Batch 3 | 424 | 133 |
| **Total** | **1,856** | **374** |

**Supplementary Table 9. Cohort-specific VCF extraction benchmark (chr22, 5 cohorts).**

| Cohort ($N$ samples) | GraphMana (s) | bcftools (s) |
|---|---|---|
| AFR (893) | 191 | 59 |
| EUR (633) | 211 | 51 |
| EAS (585) | 131 | 49 |
| EUR+EAS (1,218) | 238 | 64 |
| ALL (3,202) | 571 | 110 |

**Supplementary Table 10. Multi-format export benchmark (chr22, from single database).**

| Format | GraphMana (s) | bcftools (s) |
|---|---|---|
| VCF | 473 | 96 |
| TreeMix | 189 | N/A |
| SFS (dadi) | 86 | N/A |
| SFS (fsc) | 119 | N/A |
| BED | 84 | N/A |
| TSV | 85 | N/A |

**Annotation update** (53,000 BED regions): GraphMana **3.5 s**, bcftools 96 s (**27**×). **Lifecycle**: GraphMana 98 min / **46 ops**; bcftools 17 min / 17 ops (9 N/A).

**Supplementary Table 11. Whole-genome export timings (70.7M variants, 3,202 human WGS samples).**

| Format | Path | Variants | Wall time | Size |
|---|---|---|---|---|
| TreeMix | FAST | 70,692,015 | 102 min | 780 MB |
| SFS dadi | FAST | 70,692,015 | 98 min | 5.7 KB |
| SFS fsc | FAST | 70,692,015 | 101 min | 1.3 KB |
| BED | FAST | 70,692,007 | 103 min | 2.9 GB |
| TSV | FAST | 70,692,007 | 101 min | 3.8 GB |
| PLINK 1.9 (8 thr) | FULL | 9,627,636 | 156 s | 7.2 GB |
| VCF (BGZF) | FULL | 68,912,619 | 3.7 hr | 14 GB |
| EIGENSTRAT | FULL | 61,599,149 | 3.7 hr | 184 GB |
